# A data mining paradigm for identifying key factors in biological processes using gene expression data

**DOI:** 10.1101/327478

**Authors:** Jin Li, Le Zheng, Akihiko Uchiyama, Lianghua Bin, Theodora M. Mauro, Peter M. Elias, Tadeusz Pawelczyk, Monika Sakowicz-Burkiewicz, Magdalena Trzeciak, Donald Y. M. Leung, Maria I. Morasso, Peng Yu

## Abstract

A large volume of biological data is being generated for studying mechanisms of various biological processes. These precious data enable large-scale computational analyses to gain biological insights. However, it remains a challenge to mine the data efficiently for knowledge discovery. The heterogeneity of these data makes it difficult to consistently integrate them, slowing down the process of biological discovery. We introduce a data processing paradigm to identify key factors in biological processes via systematic collection of gene expression datasets, primary analysis of data, and evaluation of consistent signals. To demonstrate its effectiveness, our paradigm was applied to epidermal development and identified many genes that play a potential role in this process. Besides the known epidermal development genes, a substantial proportion of the identified genes are still not supported by gain- or loss-of-function studies, yielding many novel genes for future studies. Among them, we selected a top gene for loss-of-function experimental validation and confirmed its function in epidermal differentiation, proving the ability of this paradigm to identify new factors in biological processes. In addition, this paradigm revealed many key genes in cold-induced thermogenesis using data from cold-challenged tissues, demonstrating its generalizability. This paradigm can lead to fruitful results for studying molecular mechanisms in an era of explosive accumulation of publicly available biological data.

## Introduction

The huge amount of data generated from previous biological studies provides a precious resource for mining new biological knowledge. A significant portion of the data is freely available in public repositories such as ArrayExpress ^1^ and Gene Expression Omnibus (GEO) ^2^. For example, around one million series studies are publicly available in GEO. Due to the unstructured nature of the metadata associated with public data, manual curation is required ^3-7^, a step that is essential for collecting large-scale gene expression data.

Gene expression data facilitate the application of the network reconstruction approach for identifying key factors in biological processes. For example, Bhaduri et al.^8^ applied the gene network reconstruction approach to explore epidermal differentiation regulators. Using network analysis, the *MPZL3* gene was identified as a highly connected hub required for epidermal differentiation. In addition, the *MPZL3* gene indirectly regulates epidermis genes, including *ZNF750, TP63, KLF4*, and *RCOR1*, through the *FDXR* gene and reactive oxygen species.

Complementing data analyses with more relevant data improves the identification of key factors in biological processes. Even though massive expression data can provide essential insights in revealing genetic interactions, there are confounding factors or “noise” introduced by technical variations, such as batch effects ^9^. To obviate the “noise” and generate a consistent result, one solution is integrative analysis by comparing large-scale datasets ^10^. In this report, we introduce a paradigm to integrate data collection and data analysis for mining key factors in specific biological processes (**Figure 1**). To demonstrate the power of our data processing paradigm, we evaluate key factors of two applications in skin biology and energy homeostasis.

**Fig. 1.**
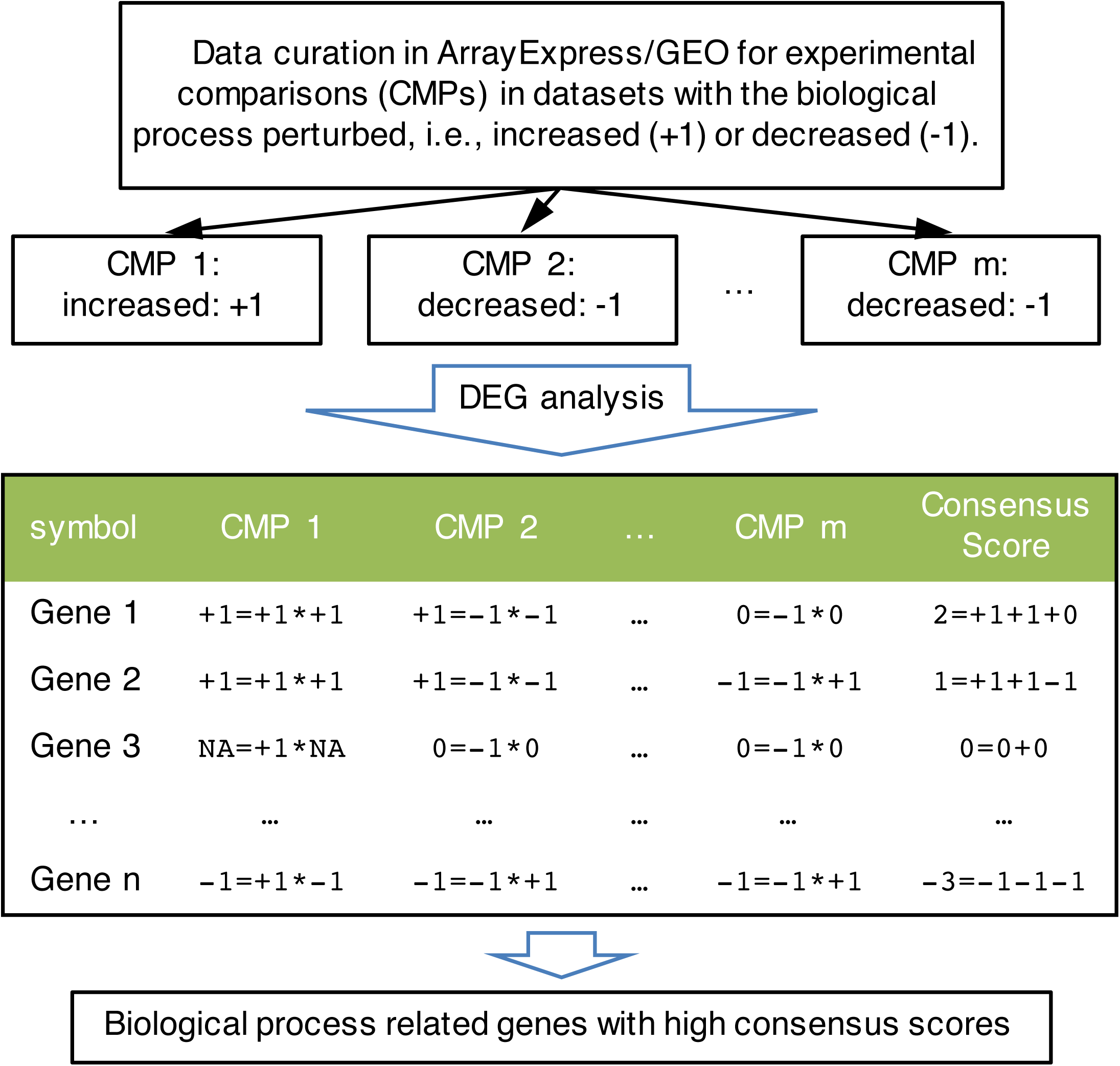
Data processing paradigm flowchart. Data curation was performed to identify the gene expression datasets with the given biological process perturbed (e.g., the process is increased in CMP 1 with +1 and is decreased in CMP 2 or CMP m with direction −1). DEG analysis was performed on the curated datasets, and +1/−1/0 represents the up-regulated, down-regulated, or unchanged genes, respectively. To prioritize important genes in the biological process for each gene in a curation dataset, an affinity score of +1/−1/0 was calculated first by comparing the gene expression change and the regulation of the biological process, where +1 indicates that the gene (e.g., Gene 1 in CMP 1 and CMP 2) is positively related to the biological process, −1 indicates that the gene (e.g., Gene 2 in CMP m and Gene n in CMP 1) is negatively related to the biological process, and 0 indicates no relation of the gene to the biological process. No measurement (notated as NA, e.g., Gene 3 in CMP 1) indicates an unknown affinity of the gene in the dataset. By summing the affinity scores, a consensus score was calculated for genes in the perturbed datasets. Genes with higher consensus scores were identified as more related to the biological process.

The epidermis of skin mediates various functions that protect against the environment, such as microbial pathogen challenges, oxidant stress, ultraviolet light, chemicals, and mechanical insults ^11^. Therefore, it is critical to understand mechanisms of epidermal development to develop new treatment for human skin diseases ^12^. Our paradigm predicts key factors in epidermal development by collecting related datasets and integrating the information. A fraction of genes are annotated in Gene Ontology (GO) or have strong functional validation based on gain-/loss-of-function studies ^13^. The remaining genes are novel; their functionality has not been experimentally validated. We picked a top hit, suprabasin (*SBSN*), and performed loss-of-function experiments for the mouse homolog of gene *Sbsn* using RNA-Seq. The analysis validates that *Sbsn* knockdown in mouse keratinocyte cultures down-regulates cornified envelope genes, suggesting an essential role of *SBSN* in epidermal differentiation. These results demonstrate the effectiveness of our paradigm in discovering key factors of epidermal development.

As another application, cold-induced thermogenesis (CIT) can reduce body weight by increasing resting energy expenditure in mammals ^14^. Genes involved in CIT can be promising therapeutic targets for treating obesity and diabetes. Thus, it is important to understand the underlying mechanism of CIT. Our paradigm detected potential CIT-related genes, including known CIT genes and novel ones, showing that the paradigm can be generalized easily to other biological processes. It is a promising integrative analysis approach to identify key factors in biological processes.

## Results

### Identification of candidate epidermal development genes

To identify key gene expression datasets that are likely to be related to epidermal development, data curation was performed. A total of 295 epidermis development genes (according to GO) were searched on ArrayExpress to query microarray datasets, and over 300 datasets were retrieved. Due to the limitation of the search function in ArrayExpress, many retrieved datasets did not have any perturbation of these epidermis development genes, even though the gene symbols were mentioned in the datasets. To overcome this problem, manual curation was performed on each retrieved dataset to retain relevant ones, and the manual curation resulted in 24 experimental comparisons from 17 datasets with gain or loss function of 14 epidermis development genes (**Table S1** and **Methods**).

To determine the candidate genes potentially involved in epidermal development, differential gene expression (DEG) analysis was performed on the 24 experimental comparisons of the curated microarray datasets. Differentially expressed genes were identified under q ≤ 0.05. The large-scale gene expression changes derived from our curated datasets provided a list of candidate genes that may be potentially involved in epidermal development (**Figure 2**).

**Fig. 2.**
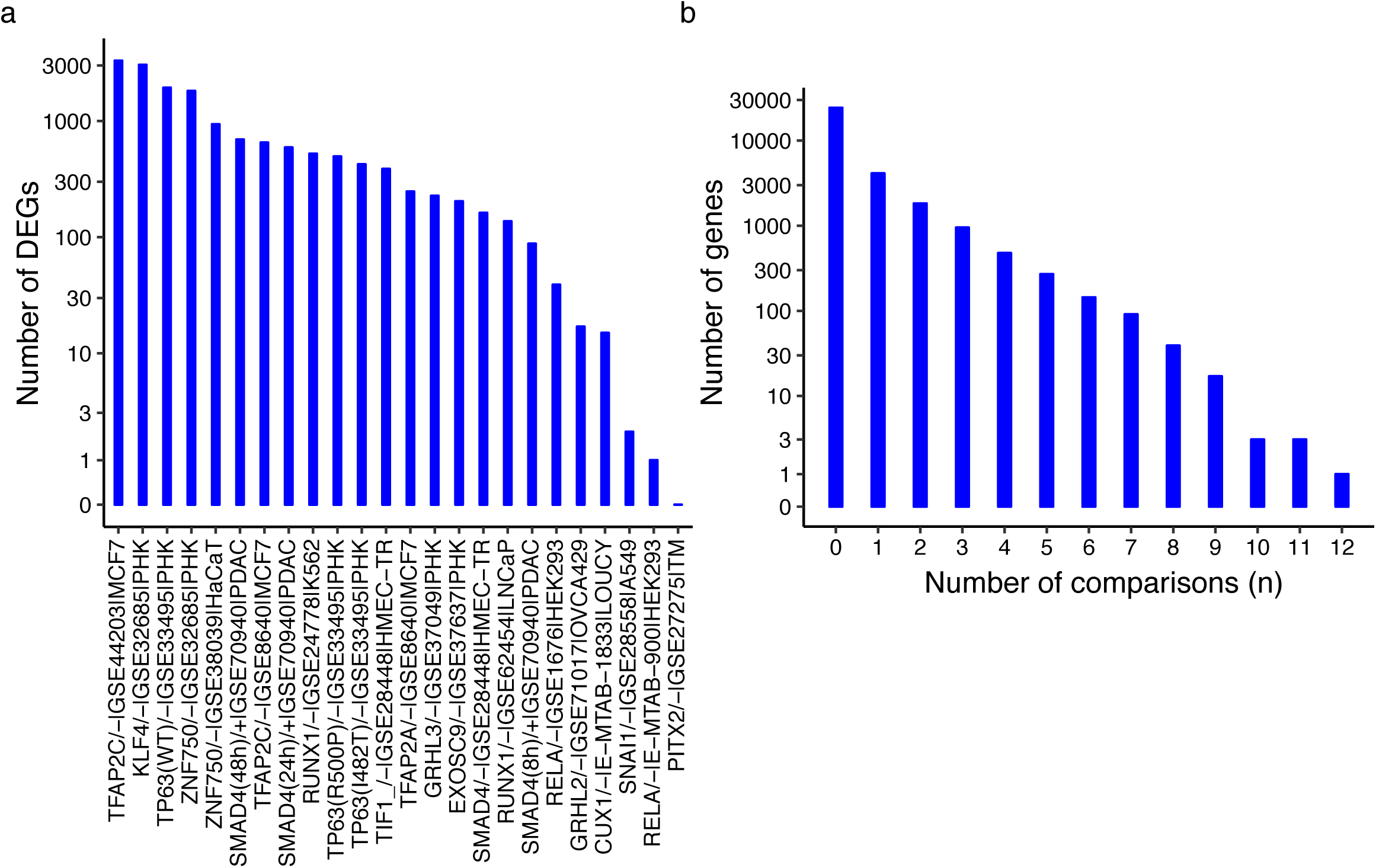
DEG results of the curated microarray datasets. To identify the differentially expressed genes of the 24 experimental comparisons in curated microarray datasets, DEG analysis was performed as mentioned in supplemental materials. DEGs were identified under q ≤ 0.05. (a) The bar plot depicts the number of DEGs identified in each of the 24 experimental comparisons. (b) The figure depicts the number of genes differentially expressed in *n* comparisons out of all the 24 comparisons. A large number of DEGs were identified in the curated datasets. A small group of genes was differentially expressed in multiple datasets.

To identify genes that are potentially critical in epidermal development, consensus gene scores were summarized for each gene from affinities on the 24 experimental comparisons. Eighty-one genes were identified as key genes related to epidermal development with a consensus score ≥ 6 (**Table S2**). The heatmap (**Figure S1**) shows a majority of these genes with a +1 affinity score in skin-related cell types.

This information suggests that these top genes may play a role in epidermal development. To infer the biological processes involved, GO analysis was performed on these top genes using Fisher’s exact test (the null hypothesis is log-odds-ratio < 2) with all the genes annotated in GO as the background. Several epidermis-related GO terms were enriched in these genes (**Figure 3**). For example, the essential GO terms in the epidermis were enriched, such as keratinocyte differentiation, epidermal cell differentiation, epidermis development, skin development, cornified envelope, and keratinization. In addition, the GO terms involved in skin barrier formation were also enriched, such as fatty acid elongase activity, lipoxygenase pathway, and establishment of skin barrier. These enriched GO terms suggest that the top identified genes are critical in epidermal development.

**Fig. 3.**
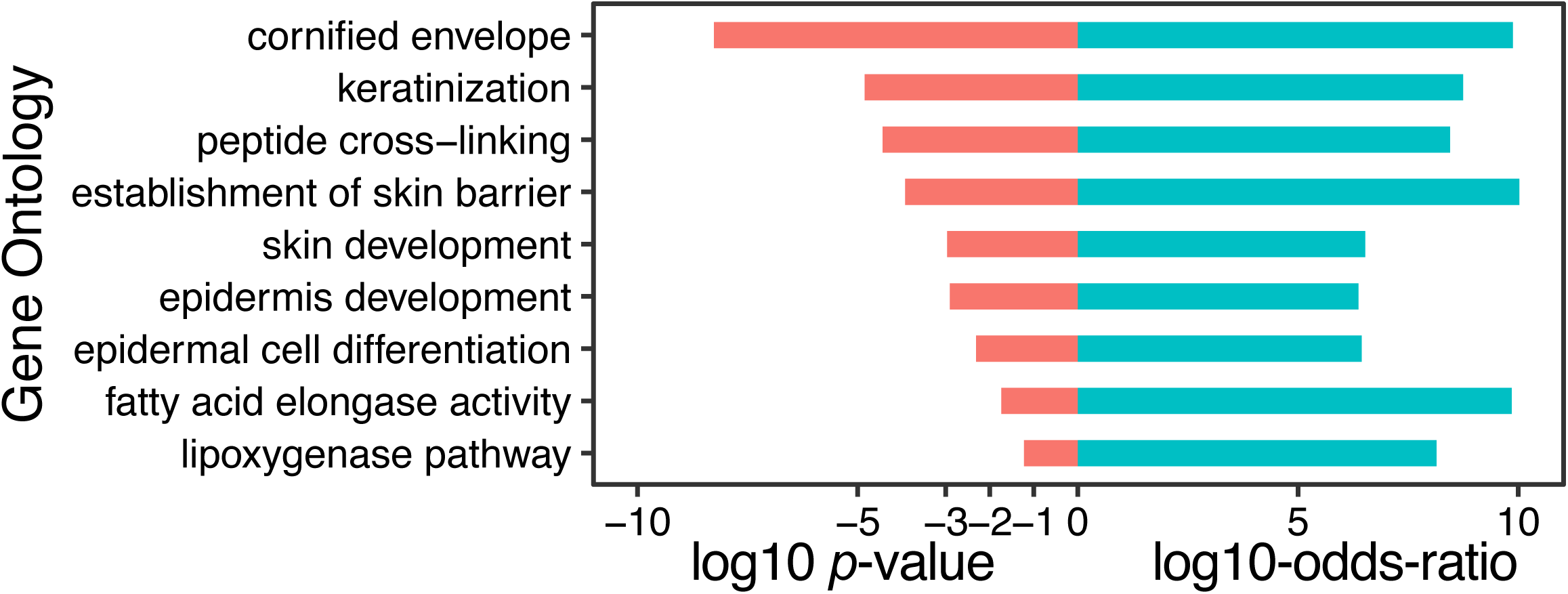
Biological process and literature study of genes with consensus score ≥ 6. To identify the biological process that the 81 top genes (consensus score ≥ 6) were involved in, a GO enrichment analysis was performed. The enriched GO pathways were plotted with a log10 *p*-value, along with their log10 odds ratios in the enrichment analysis.

Because GO annotation is not complete for gene functions ^15^, we manually curated functional annotations for the top identified genes. Of these genes, besides the 18 genes annotated in the GO term “epidermis development,” only three genes have loss-of-function experiments supporting their role in epidermal development. However, the majority of these identified genes have no functional experimental validation on epidermal development. Of the three genes with literature evidence, *EDN1* (consensus score = 7) mediates the homeostasis of melanocyte (located at the bottom of epidermis) *in vivo* upon ultraviolet irradiation ^16^. The loss function of *ELOVL4* (consensus score = 6) represses the generation of very-long-chain fatty acids, which is critical for the epidermal barrier function, showing the important role of *ELOVL4* in epidermis development ^17^. The *in vitro* loss-of-function experiment of *HOPX* (consensus score = 6) leads to increased expression of cell differentiation markers in human keratinocytes, demonstrating its involvement in epidermal development ^18^.

To evaluate how well the roles of the identified genes are understood in epidermal development, we queried the PubMed literature database and examined the results. For each gene, the keyword used in the PubMed search was constructed as “<symbol>[tiab] AND (epidermis OR skin)”. The search results showed that a large proportion of identified genes (∼42% = 34/81) have no publications related to skin. Therefore, these understudied novel genes revealed potential candidate genes for new studies on epidermal development. In addition, the majority (> 70%) of identified genes were not in the epidermis development GO term (**Figure S2**). These novel genes demonstrate the ability of the paradigm to discover unknown factors in epidermal development.

To demonstrate the effectiveness of the paradigm computationally, top-ranked genes using collective comparisons were compared to genes using individual comparisons (**Text S1**). **Figure S3** shows the significantly (*p*-value = 3.6 × 10^−9^) increased epidermal development genes identified by the paradigm compared to differentially expressed genes derived from individual comparisons.

### Validation of Sbsn role in epidermal differentiation by loss-of-function and other experiments

Among the identified genes, a top gene (*SBSN*) (with a high consensus score of 9) was selected to validate its role in epidermal development. A phylogenetics-based GO analysis revealed enriched GO terms related to epidermal development using co-evolved genes of *SBSN* (**Text S2, Figure S4**). In addition, a time-course microarray dataset showed an increased expression of *SBSN* upon epidermal differentiation (**Text S3, Figure S5**). These results suggest a potentially critical role of *SBSN* in epidermal development. To determine the cellular component that *Sbsn* is involved with, we performed a study of the differentially expressed genes in differentiating mouse primary keratinocyte cultures from mice with *Sbsn* knockdown. In *Sbsn* knockdown mouse cultures, 326 genes were up-regulated, and 161 genes were down-regulated (**Methods, Table S3, Figure 4a**). To investigate the functional roles of *Sbsn*, these differentially expressed genes were used to search for enriched GO terms ^19^ using Fisher’s exact test (null hypothesized log-odds-ratio < 2) with the genes expressed in the *Sbsn* knockdown mouse culture and the controls as background. Specifically, the cornified envelope GO term was found enriched in the genes down-regulated upon *Sbsn* knockdown (*p*-value < 0.05), and eight cornified envelope genes were down-regulated (**Table S4**). These results suggest the role that *Sbsn* may play in epidermal differentiation and cornified envelope formation.

**Fig. 4.**
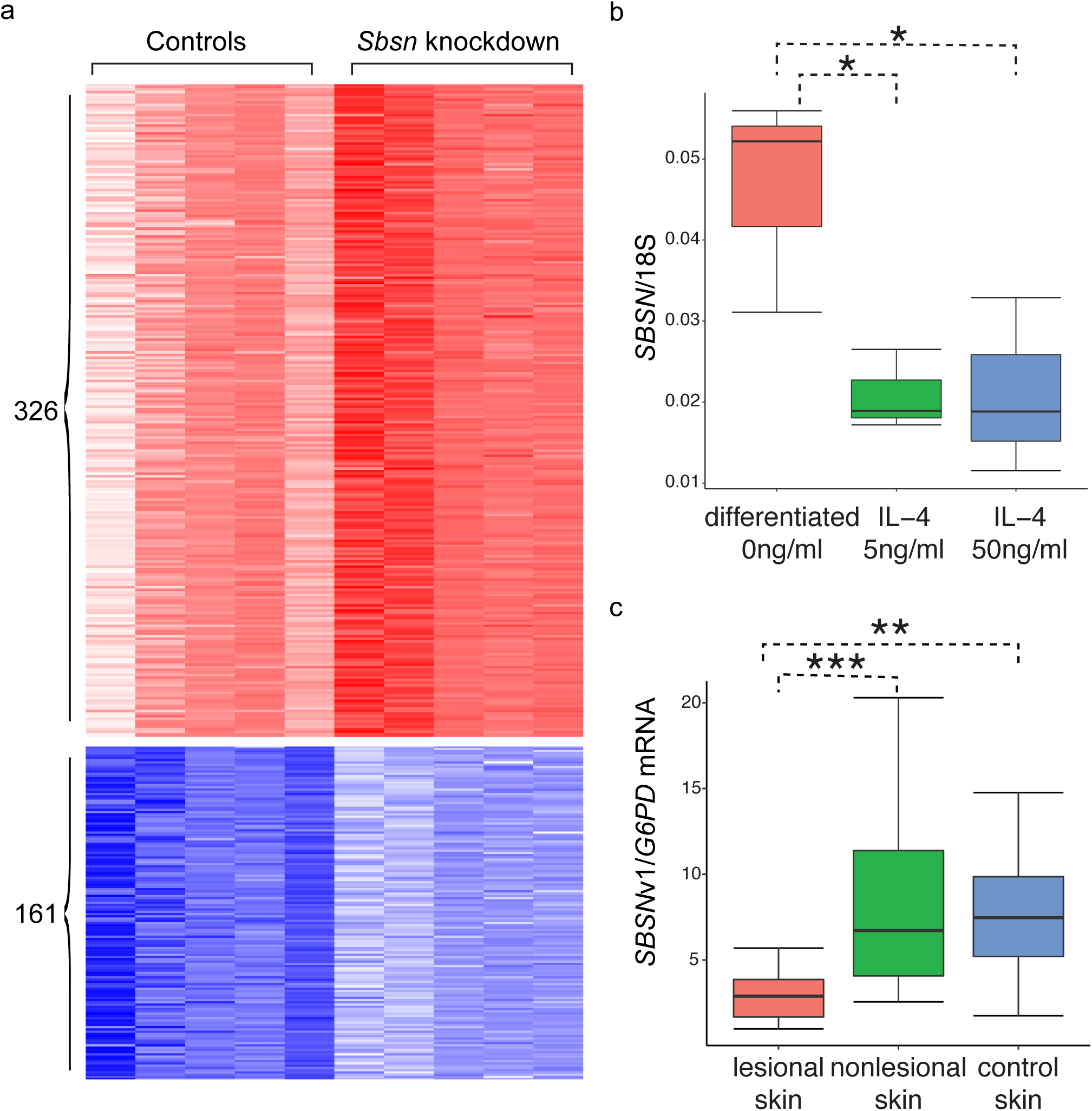
Validations of *SBSN* in epidermal differentiation. (a) Heatmap of the expression levels between *Sbsn* knockdown mice and controls. Expression levels are shown for genes differentially expressed (under |log2-fold-change| > 0.5 and *q*-value < 0.05) upon *Sbsn* knockdown. Red and white colors indicate high and low expression levels (arc-sine hyperbolic transformed normalized counts by DESeq and scaled by standard deviations) for 326 up-regulated genes, respectively. Blue and white colors indicate high and low expression levels for 161 down-regulated genes, respectively. (b) Expression values of *SBSN* normalized by 18S rRNA in differentiated keratinocytes upon IL-4 treatment. To evaluate the gene expression changes of *SBSN* during keratinocyte differentiation upon IL-4 treatment, an RT-PCR experiment was performed with nine differentiated cells with and without IL-4 treatments (three replicates per condition). The expression values of *SBSN* were normalized by the expression levels of 18S rRNA. The boxplot shows a significant decrease of *SBSN* expression at two IL doses (5 ng/ml and 50 ng/ml) (*: *p*-value < 0.05). (c) Expression values of full-length *SBSN* transcript (v1) in AD skins. To evaluate the expression changes of *SBSN* in AD skins, expression values were measured in AD skins for the *SBSN* transcripts via RT-PCR. The expression levels were normalized by the expression levels of *G6PD*. The full-length *SBSN* transcript showed significantly decreased expression levels in AD lesional skins compared to AD nonlesional and control skins (***: *p*-value < 0.001, **: *p*- value < 0.01, *: *p*-value < 0.05).

Atopic dermatitis (AD) is the most common chronic inflammatory skin disease ^20^. IL-4, a type 2 cytokine, contributes to the development of AD. Because broad defects of cornified envelope have been identified in AD ^21^, *SBSN* may play a critical role in AD via defective cornification. To investigate the putative role of *SBSN* in AD, differentiated primary normal human epidermal keratinocytes (NHEKs) were cultured to examine the expression levels of *SBSN* upon IL-4 treatments via RT-PCR. In the presence of IL-4 (at doses of 5 ng/ml and 50 ng/ml), *SBSN* mRNA levels in the differentiated cells were significantly decreased as compared to differentiated cells without cytokine treatment (**Figure 4b**). These decreased expression levels of *SBSN* upon IL-4 treatment suggest a critical precursor role of *SBSN* in the development of AD via disruption of cornification—and further indicate an important role of *SBSN* in epidermal differentiation.

To investigate the role of *SBSN* in AD, expression levels of three *SBSN* transcripts were measured in AD lesional/nonlesional and control skins via RT-PCR (**Methods**). A total of 49 skin biopsies were measured, consisting of 16 AD lesional skin biopsies, 16 AD nonlesional skin biopsies, and 17 healthy controls. The expression levels of *SBSN* transcripts were normalized to *G6PD*. *SBSN* transcript v1 (NM_001166034.1) showed a significantly decreased level in AD lesional skin compared to AD nonlesional skin and controls (**Figure 4c**). The decreased expression levels of the full-length transcript of *SBSN* suggests an important role of this *SBSN* isoform in AD.

### Generalization of the paradigm as demonstrated by its application on CIT

To investigate the generalizability of our integrative analysis approach, we applied the paradigm to reveal thermogenesis genes in tissues upon cold exposure. We collected ten gene expression datasets from GEO (**Table S5**). These gene expression data were collected from tissues of mice treated with cold temperature to induce thermogenesis. Both microarray and RNA-Seq data were collected. Because thermogenesis is always activated upon cold exposure, the direction of thermogenesis is thus increased in all the 24 comparisons within the ten collected datasets. Using DEG analysis, the paradigm calculated the consensus scores for measured genes from 24 comparisons and identified 153 genes with a consensus score ≥ 6 (**Table S6**). These 153 identified genes were then used to perform GO analysis. Enriched GO terms are related to energy homeostasis (**Figure S6**). Literature curation confirmed the functional evidence in CIT of some identified genes. For example, elongation of very-long-chain fatty acids (*Elovl3*, consensus score = 13) in ablated mice showed a proliferated metabolic rate in a cold environment, indicating a higher capacity for brown fat-mediated nonshivering thermogenesis. Thus, *Elovl3* is a key regulator for CIT in adipose tissue upon cold exposure ^22^. As another example, carnitine palmitoyltransferase 2 (*Cpt2*, consensus score = 11) depletion mediates the fatty acid oxidation in adipose tissue, which is required for CITs, suggesting the critical role of *Cpt2* in CIT ^15,23^. This second application of our paradigm in CIT suggests that the paradigm can be generalized to other biological processes. Our paradigm is a simple but important integrative data processing approach for gene expression data.

## Discussion

We propose a gene expression data processing paradigm to identify key factors in biological processes. The collection of gene expression data enhances the identification of key factors in biological processes. The application of the paradigm for epidermal development revealed known and novel epidermal development genes. To validate the novel predictions, an understudied gene, *SBSN*, was specifically investigated for its potential role in epidermal differentiation. *SBSN* has been identified in the suprabasal layers of the epithelia in the epidermis ^24^. Although *SBSN* was previously shown to be induced upon differentiation of epidermal keratinocytes, no loss-of-function study has been performed to demonstrate the functional role of *SBSN* in epidermis or skin. Our phylogenetics-based GO analysis suggests relevant biological processes in epidermal development for *SBSN* co-evolved genes (**Figure S4**). RNA-Seq analysis in *Sbsn* knockdown mouse keratinocyte cultures revealed down-regulated cornified envelope genes, suggesting a role for *Sbsn* in epidermal differentiation. *SBSN* may also be critical in AD, an inflammatory skin disease, because AD has a broadly defective cornified envelope ^21,25^. Due to the full-length isoform of *SBSN* potentially playing a more critical role in the development of AD (**Text S4, Figure 4c** and **S7**) and IL-4 being involved in AD ^20^, we examined the effects of IL-4 on human differentiating keratinocyte cultures and found decreased expression levels of *SBSN* in IL-4 treated compared to non-treated cultures (**Figure 4b**). These results indicate that *SBSN* may be a target for aberrant cytokine production in AD.

The paradigm identified key genes by collecting multiple datasets and integrating information from the collected datasets, especially from datasets of the cell or tissue types most relevant to the target biological process. However, due to the limited availability of such datasets in certain areas, relevant datasets from other cell or tissue types may also be used, for they generally will not worsen the results. This is consistent with the idea of ensemble learning, in which merging many weak and independent classifiers will result in a strong classifier ^26^. The heatmap of identified genes (consensus score ≥ 6) showed seven experimental comparisons from epidermal cells clustered together (**Figure S1**). To systematically cluster the 24 experimental comparisons, a hierarchical clustering analysis using an affinity distance metric (**Text S5**) grouped the comparisons into eleven distinct clusters (at a cutoff distance ≤ 0.05) (**Figure S8**). As a result, the seven experimental comparisons from epidermal cells were also consistently clustered together in the hierarchical clustering analysis. These results indicate that the experimental comparisons from epidermal cells contributed the most. In the future, with more datasets from epidermal tissues/cells generated, it may not be necessary to include datasets from nonepidermal tissues/cells, as the marginal contribution from them is likely to be negligible.

The paradigm starts from gene expression datasets with the perturbation of a biological process. This data collection process is critical. As for the application in epidermal development, we searched ArrayExpress using the text of epidermal development for candidate gene expression datasets. However, none of the five retrieved datasets showed changes in epidermal development, leaving us with no data from reliance on the bare keyword search functionality offered by ArrayExpress (**Figure S9**). Because known genes annotated in the epidermis development GO term provide candidate factors responsible for the regulation of epidermal development, datasets with these perturbed known genes can be a starting point for our paradigm. However, it should be understood that the paradigm is not limited to a single GO term. Due to incomplete annotation in GO ^5^, genes in other GO terms, such as keratinocyte differentiation (GO:0030216) and epidermal cell differentiation (GO:0009913), can also play roles in epidermis development. Thus, starting from these additional genes along with the genes in the epidermis development GO term, the performance of the paradigm is expected to improve because more information may be borrowed from other relevant datasets. To apply our paradigm, it is critical to examine the collection of gene expression datasets. In addition, our paradigm can include both microarray and RNA-Seq data, as shown in the CIT application, enabling the inclusion of more data—leading to better results than with only one data type.

To obtain a manageable number of identified genes, our computation analysis focused on genes with a consensus score ≥ 6. A simulation was performed to evaluate the empirical distribution of consensus scores in epidermal development and CIT, and a cutoff of ≥ 6 corresponds to an empirical *p*-value = 3.72 × 10^−8^ and 1.09 × 10^−8^for epidermal development and CIT, respectively (**Text S6** and **Figure S10**). But other thresholds may also be used. Higher thresholds lead to fewer but more robust identified genes, while lower thresholds lead to more but less robust identified genes. An investigator should pick a cutoff appropriate for the intent of the investigation. For example, if the purpose is to identify more novel epidermis development genes for further experimental validation, genes with consensus scores lower than 6 can also be considered. In addition, the cutoff should be also related to the total number of comparisons used in the analysis. In general, with more comparisons, the cutoff for the consensus score should be greater.

It is worth mentioning that genes with negative consensus scores, in general, do not contribute to regulation in biological processes. For the application of epidermal development and CIT, a consensus score cutoff ≤-5 is used to extract negative genes, and no enriched GO terms are related to epidermal development (44 negative genes) and CIT (95 negative genes) (data not shown). These results are consistent with the intent of the scoring scheme defined in **Figure 1**. The number of genes with positive and negative consensus scores would be expected to be roughly the same due to normalization, but the positive genes, in fact, have longer tails than the negative genes (**Figure S11**). Quantile testing shows significantly larger positive scores compared to the absolute negative score at 0.95-quantile for both applications (*p*-value < 2.2 × 10^−16^) ^27^. This suggests that genes positively correlated with epidermal development and CIT are more likely to be consistent across different experiments, indicating that these positive genes are more likely to be relevant to the respective processes. In summary, the paradigm is valuable in identifying key factors for biological processes using gene expression data.

## Methods

### Ethics statement

All skin samples were collected according to procedures (NKEBN/486/2011) previously approved by the local ethics committee (Independent Bioethics Commission for Research at Medical University of Gdansk). Written consent was obtained from all patients prior to enrollment in the study.

### Curating gene expression data related to epidermal development

We collected gene expression datasets related to epidermal development by manual curation according to the following procedure. First, we searched ArrayExpress using the keyword (“epidermis+development” OR “epidermal+development”) AND organism: “homo sapiens”, retrieving only five studies, none of which could be reused to study the epidermal development process because of no change in epidermal development in the datasets (**Figure S9**). Therefore, we started from known epidermal development genes to curate datasets with the process perturbed. Specifically, genes from the GO ^19^ epidermis development (accession GO:0008544) term were extracted first for humans. Then, the official symbol of each gene was queried on ArrayExpress for human microarray datasets. Each retrieved dataset was manually examined to retain only the datasets with at least one epidermis development gene being perturbed (i.e., knocked out, knocked down, or overexpressed). To ensure proper downstream statistical analysis, any dataset with no replicates was discarded.

### Data processing paradigm of the perturbed expression data

To identify the genes related to a biological process, our data processing paradigm was performed on the gene expression data to capture the affinities between specific genes and the biological process. An affinity score of +1 or −1 means that the gene is positively or negatively related to the biological process. Specifically, if the expression of a gene is increased or decreased in a biological process that is increased, the gene has an affinity score of +1 or −1 for the biological process. Alternatively, if the biological process is decreased, these genes have an affinity score of −1 or +1. The affinity score was 0 or NA for the genes not differentially expressed or unmeasured. The detailed workflow of the paradigm is shown in **Figure 1**. For a biological process, systematic data curation is performed to collect gene expression datasets with the process perturbed (increased or decreased). Using DEG analysis (**Text S7** and **S8**) ^28-30^, affinity scores are calculated for each gene in each comparison in each dataset. Finally, a consensus score is calculated by summing these affinity scores among the comparisons for each gene. High consensus scores suggest that the corresponding genes are potentially critical to the biological process. Thus, our paradigm is a general framework that can be used to identify the key factors in a biological process.

### NHEK cell culture and treatment

Primary NHEKs of neonatal foreskin were purchased from Thermo Fisher Scientific and were maintained in EpiLife Medium containing 0.06 mM CaCl_2_ and S7 supplemental reagent under standard tissue culture conditions. The cells were seeded in 24 well dishes at 2 × 10^5^/well to form a confluent monolayer. In the following day, the cells were subjected to differentiation by increasing CaCl_2_ to 1.3 mM in the culture media with or without the human recombinant IL-4 at designated concentrations. The cells were harvested for total RNA extraction before differentiation and were differentiated for 5 days. Total RNA was extracted using RNeasy mini kit according to manufacturer guidelines (QIAGEN, MD). RNA was then reverse transcribed into cDNA using superScript® III reverse transcriptase from Invitrogen (Portland, OR) and was analyzed by real-time RT-PCR using an ABI Prism 7000 sequence detector (Applied Biosystems, Foster City, CA). Primers and probes for human *SBSN* (Hs01078781_m1) and *18s* (Hs99999901_S1) were purchased from Applied Biosystems (Foster City, CA). Quantities of all target genes in test samples were normalized to the corresponding 18S levels.

### Sbsn siRNA in mouse keratinocyte culture and RNA sequencing

Primary murine keratinocytes were isolated from BALB/c neonatal mice as previously described ^31^. Primary murine keratinocytes were cultured in a supplemented minimal essential medium (Gibco, Thermo Fisher Scientific Inc., Waltham, MA, USA) with 8% fetal bovine serum (FBS, Atlanta Biologicals, Flowery Branch, GA, USA) and 1% antibiotic (Penicillin Streptomycin Amphotericin B, Sigma), with 0.05 mM Ca^2+^ concentration. A total of 1×10^6^ cells were seeded in each well of six well plates. Twenty-four hours after seeding, the siRNA mix (Opti-Mem serum free media (GIBCO), 75 pmol of siRNA (Dharmacon), and HiPerfect transfection reagent (Qiagen)) was added to cells. For *SBSN*, the siRNA used were Dharmacon, SMARTpool; siGENOME *Sbsn* siRNA M-054578-01-0005 and Control (mouse) SMARTpool; and siGENOME Non-Targeting siRNA Pool #1. Each condition was done in triplicates. To induce keratinocyte differentiation, a final concentration of 0.12 mM Ca^2+^ was used. RNA was harvested 48 hours after siRNA transfection. Total RNA from cells was extracted using RNeasy kit (Qiagen) according to manufacturer instructions. A total of 100 ng was used to prepare the libraries utilizing a Neoprep Library kit (Illumina). RNA sequencing was performed in the NIAMS Genome Core Facility at the National Institutes of Health.

### DEG analysis using Sbsn knockdown RNA-Seq data in mouse differentiating primary keratinocyte cultures

To identify the differentially expressed genes in mouse differentiating primary keratinocyte cultures in which *Sbsn* had been knocked down with siRNA, the following analysis was performed. The raw RNA-Seq reads were aligned to the mouse (mm10) genome using STAR (version 2.5.1b) ^32^ with default settings. The uniquely aligned reads were retained to calculate the read counts for each gene against the UCSC KnownGene annotation (mm10), and a count table was constructed by counting the number of reads aligned uniquely to each of the genes for each sample. DEG analysis was performed by DESeq2 ^33^. To adjust the batch effect, a generalized linear model with a batch factor was used to model the read counts for all samples, and the Wald test was used to test the significance of differences in gene expression between *Sbsn* knockdown samples and controls. FDR adjusted *q*-values were then calculated from the *p*-values in the Wald test using the Benjamini-Hochberg procedure ^34^. The log2-fold changes between *Sbsn* knockdown samples and controls were also calculated for each gene. The differentially expressed genes were identified under |log2-fold-change| > 0.5 and q < 0.05.

### RT-PCR analysis of AD skin

For the current study, arm skin samples (2 mm punch biopsies of 3 mm depth) were taken from AD patients (from lesional and nonlesional AD skin), and skin samples (controls) were obtained from healthy subjects. The nonlesional skin biopsy was performed at a 10 cm distance (at least) from AD skin lesions. Immediately after biopsy, the skin samples were placed in RNA-later solution (Qiagen) and were stored at −20°C. Total RNA was isolated using standard methods. The mRNA levels were analyzed by real-time RT-PCR with TaqMan primer-probe sets using the Path-ID Multiplex One-Step RT-PCR kit (Path-ID™ Multiplex One-Step RT-PCR Kit (Applied Biosystems). The reference transcript *G6PD* was used as an internal standard and was amplified together with each target gene transcript in the same well using primers and probes, as shown in **Table S7**. The level of each analyzed transcript was normalized to that of the appropriate reference transcript.

### Data availability

The datasets used in this study are available in the National Center for Biotechnology Information’s (NCBI’s) GEO ^2^ with accession GSE100100.

## Acknowledgements

The authors thank Yu Chen (ECE Department at Texas A&M University), Veronica Nagle (Laboratory of Skin Biology, NIAMS, NIH), and Stephen Brooks from the NIAMS Biodata Mining core facility (NIAMS, NIH) for their contributions to this work. This work was supported by startup funding to PY from the ECE department and Texas A&M Engineering Experiment Station/Dwight Look College of Engineering at Texas A&M University, by funding from TEES-AgriLife Center for Bioinformatics and Genomic Systems Engineering (CBGSE) at Texas A&M University, and by TEES seed grant. DYL was supported by NIH grants R01 AR41256 and U19 AI117673. This work was supported by IRP NIH ZIA-AR041124 (MIM), R01AR051930 (TM) and R01AG028492 (PE), which were administered by the Northern California Institute for Research and Education and with resources of the Research Service, Department of Veterans Affairs. The funders had no role in study design, data collection and analysis, decision to publish, or preparation of the manuscript.

## Author Contributions

J.L., L.Z. and P.Y. carried out the analyses and wrote the manuscript. A.U., L.B., T.M., P.E., T.P., M.SB., M.T., D.L. and M.M. performed validation experiments. All the authors reviewed and approved the final manuscript.

